# Intraspecific differences in habitat depth in a deep-sea isopod, *Bathynomus doederleini* (Crustacea: Isopoda: Cirolanidae), off the west coast of Kyushu, Japan

**DOI:** 10.1101/2025.06.13.659626

**Authors:** Sayano Anzai, Shogo Tanaka, Shimpei Tsuchida, Riko Kato, Siti Syazwani Azmi, Shoma Izumi, Yutaka Maruyama, Sota Hoshina, Satoshi Masumi, Itaru Aizawa, Jun Uchida, Tsukasa Kinoshita, Nobuhiro Yamawaki, Takashi Aoshima, Yasuhiro Morii, Kenichi Shimizu, Mitsuharu Yagi

## Abstract

The giant deep-sea isopod, *Bathynomus doederleini*, is a benthic scavenger distributed in the northwestern Pacific. Despite its ecological importance, little is known about its habitat use and intraspecific variation in body size in relation to depth. In this study, we examined the habitat depth, size structure, and distributional limits of *B. doederleini* off the western coast of Kyushu, Japan, using baited traps deployed at depths ranging from 151 to 821 m. A total of 1,152 individuals were collected, with the highest catch per unit effort (CPUE) observed between 300 and 500 m. CPUE declined sharply below 700 m, likely due to thermal constraints and interspecific competition. Body size distribution varied significantly with depth: minimum body size increased with depth, while maximum body size remained constant. Smaller individuals were more abundant in shallower, warmer waters, suggesting ontogenetic habitat segregation possibly driven by metabolic and competitive factors. No brooding individuals were captured, supporting previous findings that reproductive females avoid baited traps. These results suggest that *B. doederleini* forms a reproductively active population in the East China Sea, with ecological adaptations to thermal conditions and depth-related niche partitioning. This study highlights the importance of trap type and environmental gradients in understanding deep-sea species ecology.

## INTRODUCTION

The deep-sea isopod, genus *Bathynomus*, represents a biologically and ecologically important taxon, characterized by species diversity, wide geographic and bathymetric distribution, and extreme body size, making it a valuable model for understanding scavenging behavior and ecological adaptation in deep-sea environments. *Bathynomus* inhabits mid-depth regions of the western Pacific, ranging from 37° S to 35° N, and in the western Atlantic, from 29° S to 31° N (Lowry and Dempsey, 2006). There are currently 23 recognized species of *Bathynomus* (Ahyong., 2025). These isopods, which provide a typical example of deep-sea gigantism, rapidly locate and consume large carcasses on the deep-sea floor (Sekiguchi et al., 1982; Yagi et al., 2025), where they spend most of their time. The genus *Bathynomus* comprises mobile and active scavengers (Barradas-Ortiz et al., 2003; Sekiguchi et al., 1982). Scavengers are widely recognized for stabilizing food webs by facilitating organic matter recycling (DeVault et al., 2003). Nevertheless, ecological dynamics of deep-sea scavenging remain poorly understood, highlighting the need for further research (Beasley et al., 2012).

The relationship between body size and habitat depth in deep-sea isopods has been noted across species, but its underlying patterns, particularly at the intraspecific level, remain unclear. Regarding their habitat depth and body size, *B. obtusus* (body length (BL): approx. 10 cm) has been reported at depths of 500–550 m off the coast of Brazil (Ferreira et al., 2023), whereas *B. pelor* (BL: approx 12 cm) has been recorded at 358 m off northwestern Australia (Lowry and Dempsey, 2006; Thomson et al., 2009). Additionally, the larger *B. giganteus* (BL: 36 cm) has been reported at depths ranging from 359 to 1050 m in the Gulf of Mexico (continental slope of the Yucatán Peninsula) (Barradas-Ortiz et al., 2003). Therefore, in interspecific relationships, larger body size may be associated with a tendency to inhabit greater depths. Scavenging fish species also exhibit increased body size with greater depth (Collins et al., 2005). However, this depth–size relationship remains poorly understood at the intraspecific level, particularly among deep-sea crustaceans.

*B. doederleini* is a relatively small species (BL: approx. 12 cm) among deep-sea isopods. With increasing commercial demand for food and ornamental purposes, this species has caught the public’s attention. It inhabits deep-sea environments (250–550 m in depth) in the northwestern Pacific, including the Philippines, Taiwan, and Japan (Sekiguchi et al., 1982; Tanaka et al., 2023). Records of *B. doederleini* captured in waters around Japan include resource surveys using shrimp traps at depths of 200–450 m in Sagami Bay (Onishi et al., 1977) and trawl and dredge surveys conducted off the coast of Okinawa (Yamashita, 1979). In the East China Sea, although *B. doederleini* was captured by the Yoko-maru research vessel of the Seikai National Fisheries Research Institute (Seikai National Fisheries Research Institute, 1988, 1995), there have been few reports. Its distribution off the west coast of Kyushu, Japan, considered its northern limit in the East China Sea, remains poorly known. This region features a gently sloping continental shelf at depths of 200–400 meters (Furuhashi et al., 2010). The present study used deep-sea bait traps to clarify habitat depth, geographic distribution, and body size of *B. doederleini* off the west coast of Kyushu, Japan.

## MATERIAL AND METHODS

Seven baited trap surveys were conducted using the Nagasaki University training vessel Kakuyo-maru between December 2021 and December 2024. Baited traps were deployed at 19 stations in the East China Sea at depths ranging from 151 to 821 meters (Fig. 1). The latitude, longitude, and depth of each station are listed in Supplementary Table 1. Baited traps consisted of a pole, buoy, line, and traps, and were designed to capture deep-sea organisms according to Sumitomo and Ueta (2003). A flag and a light (Zenipika III, Zeni Lite Buoy Co., LTD., Japan) were attached to the pole to increase visibility. Two types of traps were used: two rectangular traps (0.65 m × 0.46 m × 0.23 m, mesh size 11 mm) and one cylindrical trap (0.1 m diameter × 0.4 m width, mesh size 2 mm) (Fig. 2). To increase trap weight, steel rebar was attached to trap frames. The line was 600 lb nylon and was tied to a float using a reed line. Line lengths ranged from 500 to 1500 m, depending on depth. Each trap was baited with approx. 0.5 kg of fish chunks, primarily chub mackerel (*Scomber japonicus*). Traps were sunk during the day and retrieved the next day, except during Voyage 2 (Supplementary Table S1), which occurred 30 days later due to bad weather. Water temperature at each station was acquired with a CTD (SBE-911plus, Sea-bird Science, U.S.A). In addition, a temperature data logger (model UTBi-001, Onset Computer Corporation, MA, U.S.A.) was installed in the trap to observe water temperatures of stations at depths greater than 500 m because CTD observations could not be carried out due to wire length. Trap time and water temperature of each station are also listed in Supplementary Table S1.

**Fig. 1.**
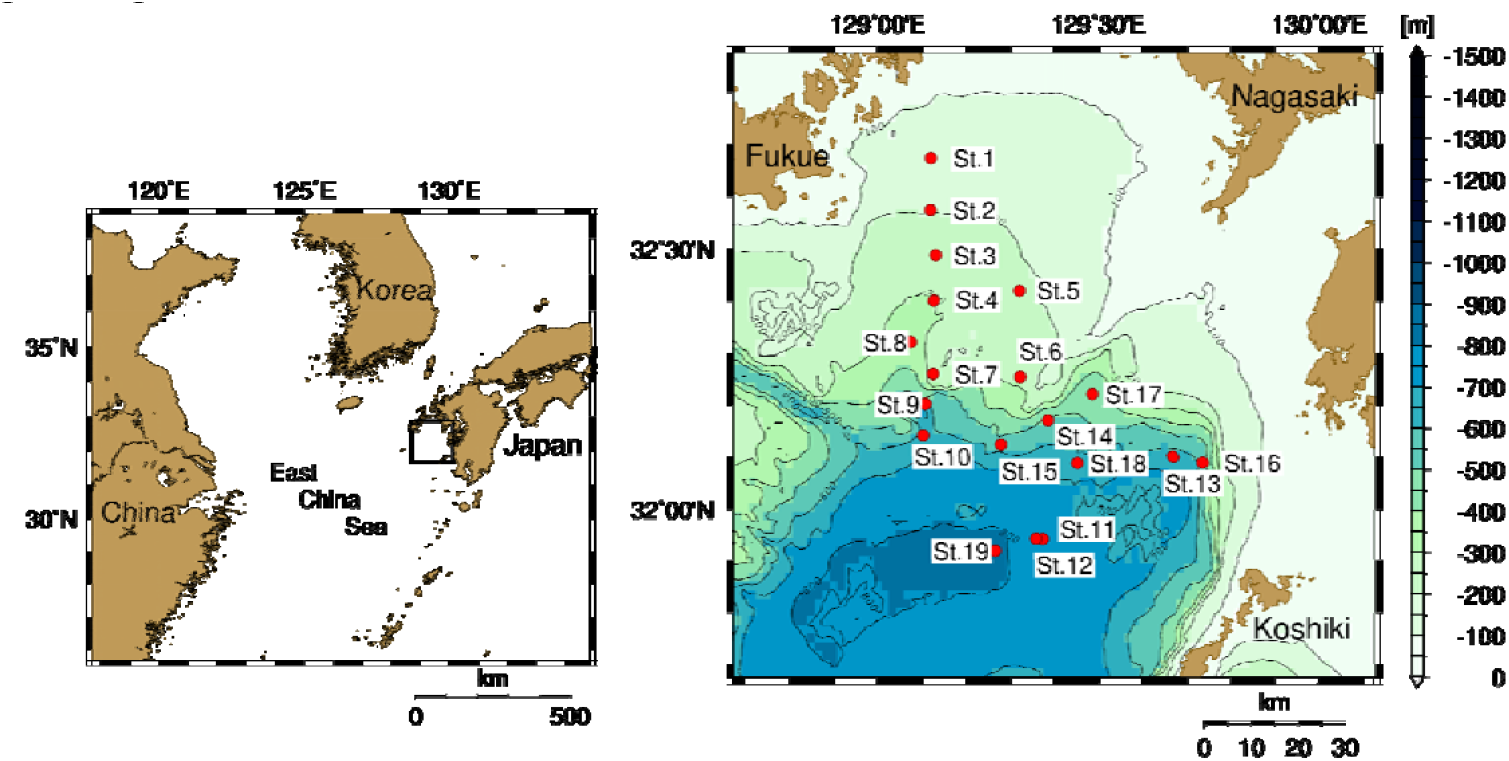
Locations of deep-sea baited trap deployed to capture *Bathynomus doederleini* in the East China Sea.

**Fig. 2.**
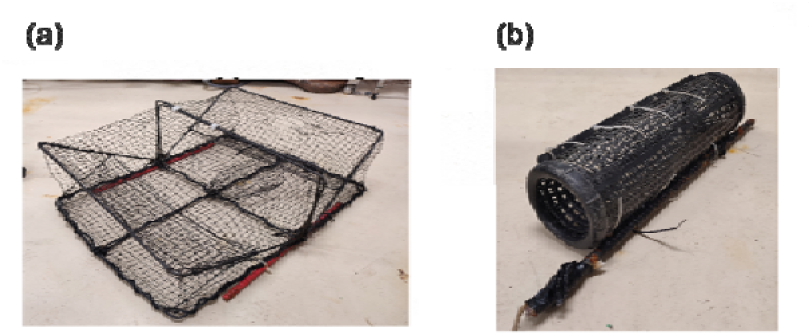
Baited traps were employed in this study. (a) A rectangular trap. (b) A cylindrical trap.

Captured organisms were brought back to the Fish and Ships Laboratory at Nagasaki University, Japan for species identification and measurements of body length and weight. Fish and decapod species were identified following Nakabo (2013), Ikeda (1998), and Miyake (1982, 1983), with some specimens identified to the family or order level. In this study, catch per unit effort (CPUE), an important indicator for assessing fishery resource abundance, was calculated by dividing the number of individuals captured by the total trap soak time and the number of traps used. However, at stations St. 3 and St. 4 during Voyage 2, traps could not be retrieved the following day due to poor weather conditions and were instead recovered 30 days later. Nevertheless, CPUE was calculated based on the assumption that the bait used to attract organisms was exhausted within 24 h (Supplementary Table 1). In practice, all bait was reduced to bones within 24 h, confirming that it no longer had any attractant effect thereafter. Additionally, data from rectangular or cylindrical traps that were found upon retrieval to have holes made by organisms, were excluded from the analysis. Statistical analyses were conducted after confirming homogeneity of variances and normality of the data. Parametric or non-parametric tests were applied depending on the results. A significance level of *p* < 0.05 was used for all statistical tests, which were performed using R (version 4.4.0).

## RESULTS

A total of 1152 *B. doederleini* were collected at depths ranging from 201 m (water temperature: 14.6 °C) to 764 m (5.7 °C) (Supplementary Table 1). In addition, other captured species included hagfish (*Eptatretus okinoseanus*) and deep-sea eels (*Simenchelys parasitica*), and the crab (*Chorilia japonica*). Detailed information is provided in Supplementary Table S2. The relationship between body length and body mass was examined on a log–log scale. It had a slope of 2.94 and an intercept of 3.54 × 10^−2^ (Fig. 3). A strong positive correlation was observed (Pearson’s correlation test: *r* = 0.98, n = 1152, *p* < 0.001).

**Fig. 3.**
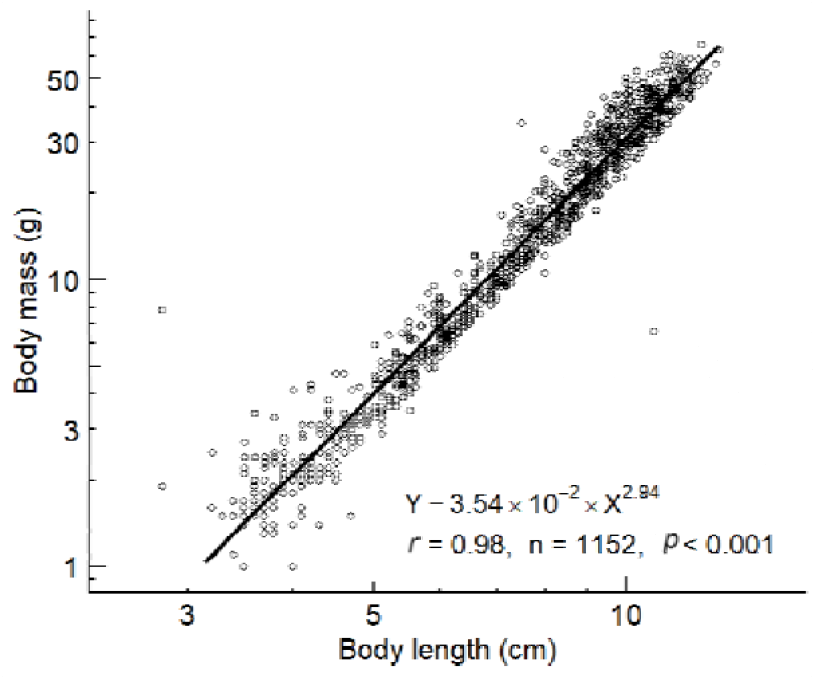
Relationship between body length and mass of the deep-sea isopod, *Bathynomus doederleini*, in the East China Sea. Both axes are logarithmic.

The highest number of individuals was recorded at St. 8 (depth: 337 m, water temperature: 9.9 °C), where 201 specimens were collected. No individuals were captured at depths of 151 m (16.7 °C) or 821 m (5.7 °C) (Supplementary Table 1). There was no significant difference in the median CPUE between trap types (Mann-Whitney U test,, n = 18 for rectangular and n = 18 for cylindrical traps, respectively; *W* = 118.5, *p* = 0.173) (Fig. 4). In contrast, when depth was categorized into 100-m intervals, a significant difference in mean CPUE among depth ranges was detected (Kruskal-Wallis test, χ^2^ (5) = 21.76, p < 0.001), with the highest CPUE observed from 400–500 m for both trap types (Fig. 5).

**Fig. 4.**
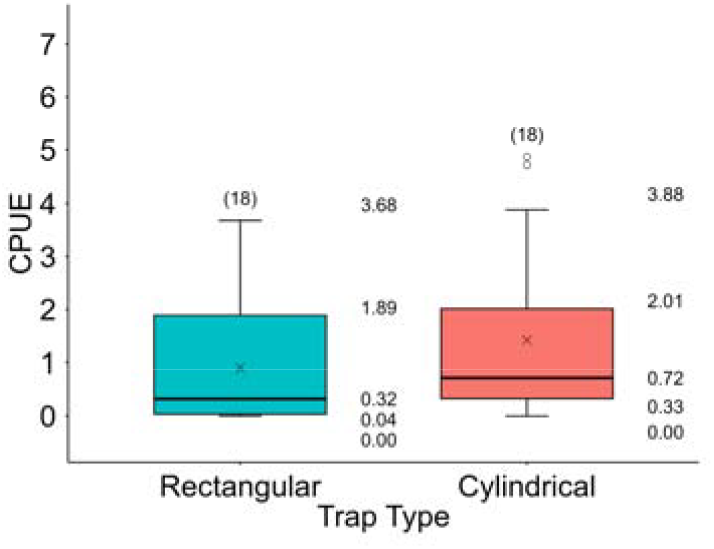
Comparison of CPUE between trap types for *Bathynomus doederleini* in the East China Sea showed no statistically significant difference in median values (Mann-Whitney U test, *W =* 118.5, *p* = 0.173). Lower and upper box boundaries indicate the 25th and 75th percentiles. X-marks indicate the means. Interior box lines are the median, and exterior whiskers are the 10th and 90th percentiles. Numbers in parentheses are sample sizes (n).

**Fig. 5.**
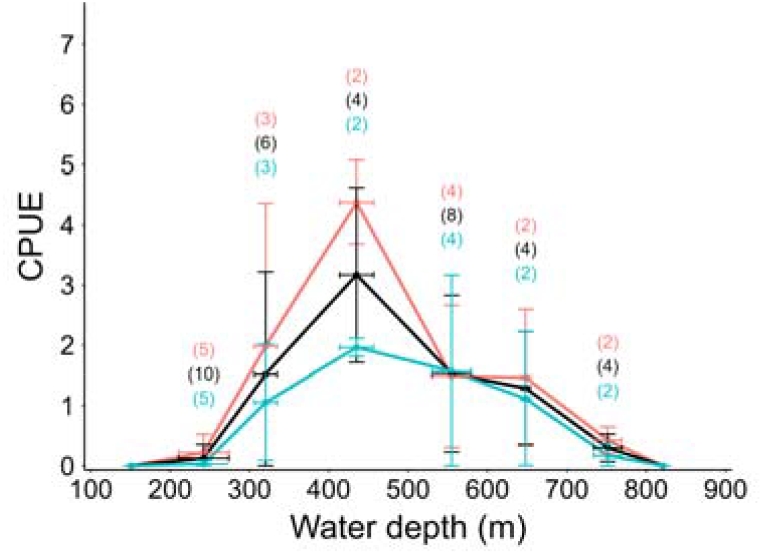
Comparison of CPUE at different depths (in 100-m intervals) for *Bathynomus doederleini* in the East China Sea. Mean capture depth and mean CPUE are shown, with error bars representing standard deviations. When depth was categorized into 100-m intervals, a significant difference in median CPUE among depth ranges was detected (Kruskal-Wallis test, χ^2^ (5) = 21.76, p < 0.001). Colors represent different trap types. Light blue indicates rectangular traps, and pink indicates cylindrical traps. Black represents the mean value of both types. Numbers in parentheses denote sample sizes (n).

The maximum body length and weight recorded were 12.9 cm and 65.9 g, respectively, while the minimum values were 2.9 cm and 0.9 g. The smallest individual was identified as a manca larva, possessing only six pairs of pereopods (Briones-Fourzán & Lozano-Alvarez, 1991). A statistically significant difference in median body size was found between trap types (Mann-Whitney *U* test, n = 626 and 506 for rectangular and cylindrical traps, respectively; *W* = 230113, *p* = 2.2 × 10^−16^) (Fig. 6). When depth was divided into 100-m intervals, a significant difference in the distribution of body length among depth ranges was observed (Kruskal-Wallis rank sum test, *H*(5)=165.56, *p* = 2.2 × 10^−16^), indicating that the size of *B. doederleini* varies with depth. Furthermore, while no significant correlation was found between mean depth and body size of the largest 5% of individuals in each depth range (Spearman rank order correlation test, n = 63, *Spearman’s* ρ = 0.00 *p* = 0.99), a significant positive correlation was detected for the smallest individuals (Spearman rank order correlation test, n = 68, *Spearman’s ρ* = 0.74, *p* = 9.5 × 10 □ ^13^) (Fig. 7).

**Fig. 6.**
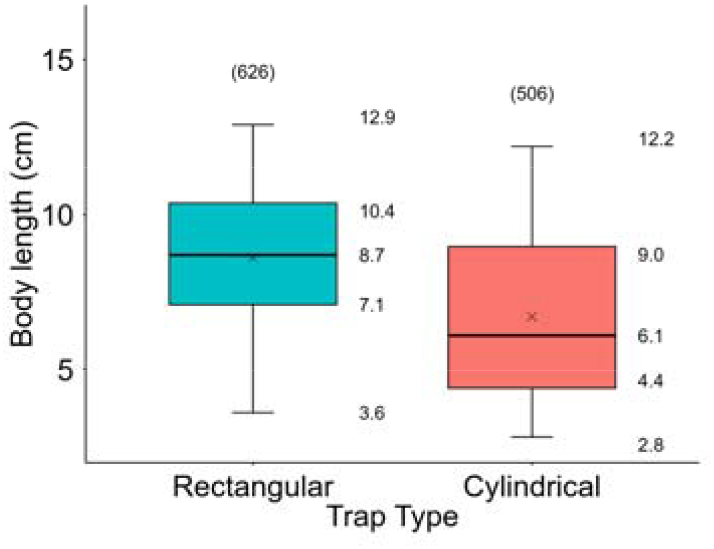
Comparison of body sizes between trap types for *Bathynomus doederleini* in the East China Sea showed a statistically significant difference in the median value (Mann-Whitney *U* test, *W* = 230113, *p* = 2.2 × 10^−16^). Lower and upper box boundaries indicate the 25th and 75th percentiles, X-marks represent the means. Interior box lines are the medians, and whiskers are the 10th and 90th percentiles. Numbers in parentheses are sample sizes (n).

**Fig. 7.**
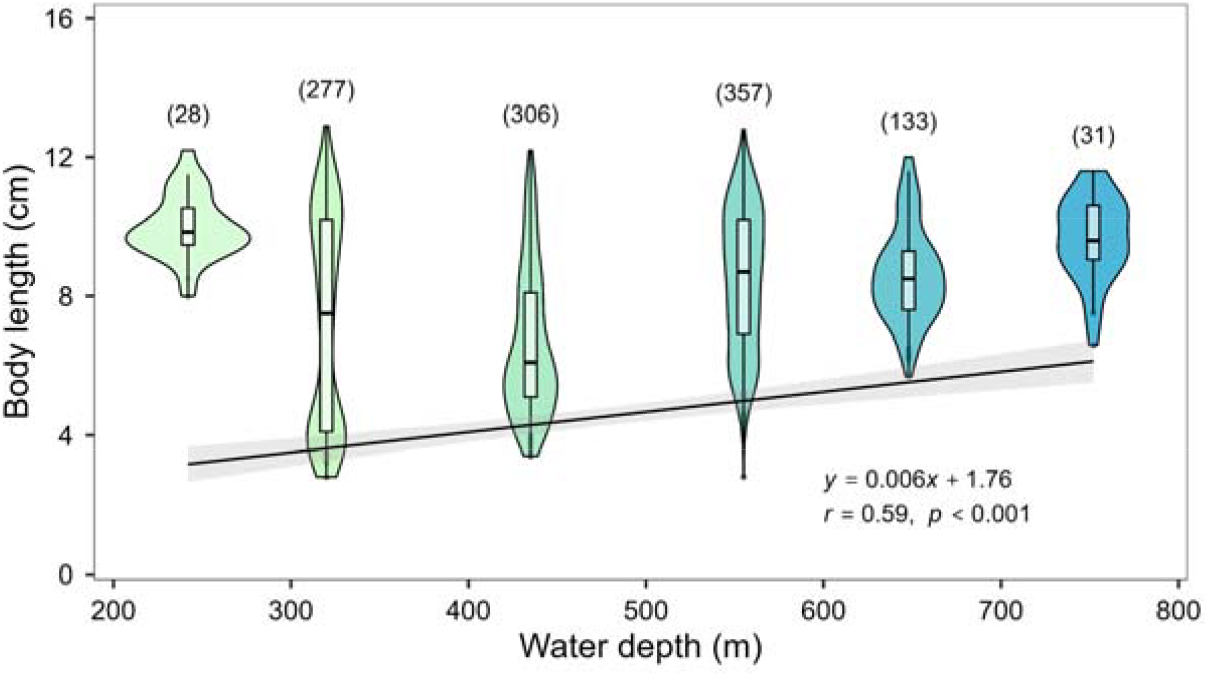
Body size distribution of *Bathynomus doederleini* in the East China Sea by depth (in 100-m intervals). Regression lines are shown for body sizes of the largest and smallest 5% of individuals in relation to mean depth. The regression for the largest 5% was not statistically significant (n = 63, Spearman’s *ρ* = 0.00 *p* = 0.99), whereas a significant relationship was found for the smallest 5% (n = 68, Spearman’s *ρ* = 0.74, *p* = 9.5× 10?13), indicating that smaller individuals tend to be absent at greater depths. Numbers in parentheses are sample sizes (n).

## DISCUSSION

This study provides new insights into the depth distribution, abundance, and population structure of *B. doederleini* in the waters off western Kyushu, highlighting the ecological dominance of this species at its northernmost distributional limit in the East China Sea. The presence of substantial biomass along western Kyushu, together with the capture of manca larvae (juvenile form), indicates a thriving reproductive population in this area. In addition, the length–weight relationship (Fig. 3) closely matched that reported by Soong and Mok (1991), confirming the consistency of the present findings with previous research. The genus *Bathynomus* is dominant in certain habitats (Shimpley et al., 2025), and our results suggest that the species similarly dominates the southwestern waters off Kyushu, Japan. Conversely, no individuals were captured at 151 m depth, and the northern part of the study area consists of shallow coastal shelf environments. No records exist for this species in the Sea of Japan (Sekiguchi et al., 1982); thus, the Goto submarine canyon off western Kyushu, Japan is considered the East China Sea’s northernmost distributional limit of *B. doederleini*.

Trap type had no significant effect on the CPUE of *B. doederleini* (Fig. 4). Depth significantly influenced the CPUE regardless of trap type, with a unimodal peak at around 400 m (Fig. 5). This pattern closely aligns with the dominant depth distribution of the species (300–500 m) reported in Sagami Bay and in waters off the Kii Peninsula, Japan (Sekiguchi et al., 1982; Tanaka et al., 1983). Similarly, the observed decline in numbers below 500 m is consistent with previous findings from Sagami Bay, Japan (Onishi et al., 1977; Tanaka et al., 1983). Among environmental factors potentially determining this vertical distribution, water temperature is likely a key driver. In the present study area, water temperature was approximately 15°C at 200 m and decreased with depth, reaching below 6°C at 800 m (Supplementary Fig. S1). *B. doederleini* has been shown to maintain a stable metabolic response, as indicated by a consistent Q□□ value, within a temperature range of 12 - 9°C (Tanaka et al., 2023), and exhibits particularly active feeding behavior around 8°C (Sekiguchi et al., 1982). In the current survey area, the depth at which water temperature was around 12 - 8°C coincided with the 300–500 m range, where CPUE also peaked, suggesting that this range represents the optimal temperature for *B. doederleini*. In contrast, CPUE declined sharply below 700 m. This may be attributed not only to limitations on feeding activity and mobility imposed by lower temperatures, but also to competitive exclusion by other scavengers sharing similar trophic niches. Indeed, several carrion-feeding fishes, such as the deep-sea eel (*Simenchelys parasitica*) and hagfish (*Eptatretus* sp.), were frequently captured at depths below 700 m. These species have been collected together with *B. doederleini* in waters around Japan (Sekiguchi et al., 1981; Sumitomo and Ueta., 2003). Similar interspecific competitive interactions among scavengers have been reported from other regions (Yeh and Drazen, 2009), indicating that assemblage composition at baited sites may be influenced by such dynamics. Moreover, strong vertical zonation in species composition and abundance has been widely documented in deep-sea environments. In the present study area, southwest of the Goto Islands, Japan, shifts in fish community structure have also been reported below 600 m (Furuhashi et al., 2010). Therefore, the observed decline in CPUE below 700 m likely reflects not only thermal constraints, but also changes in species composition associated with depth stratification.

The mean body length of *B. doederleini* captured in the cylindrical traps was significantly smaller than those captured by rectangular traps (Fig. 6). This difference likely reflects the variation in mesh size between trap types (Fig. 2). The cylindrical traps had smaller mesh openings (2.8 cm) compared to the rectangular traps (3.6 cm), allowing the capture of smaller individuals, including manca larvae. In contrast, the maximum body lengths of individuals captured by each trap type were similar, 12.2 cm for cylindrical traps and 12.9 cm for rectangular traps, indicating no clear selectivity bias for larger individuals. Therefore, cylindrical traps with smaller mesh sizes may be more suitable for studying a wide range of sizes. Moreover, a greater diversity of deep-sea organisms was incidentally captured by the rectangular traps (Supplementary Table S3), which is likely because the entrance of the cylindrical traps (Fig. 2) was more selective for species with burrowing behaviors or elongated bodies, such as *Bathynomus* and hagfish.

In this study, the body length distribution of *B. doederleini* varied across depths, showing non-uniform patterns (Fig. 7). While maximum body length was not influenced by depth, minimum body length exhibited a significant positive relationship with increasing depth, indicating that smaller individuals were absent or less abundant at greater depths. A similar depth-related size distribution has also been reported in Taiwanese waters, and it has been suggested that different life stages of *Bathynomus* may exhibit varying bait affinities (Tso and Mok, 1991). Juveniles, which require more energy for growth and maturation, are presumably more strongly attracted to bait. In the present study area, the tendency for minimum size to increase with depth may be explained by several factors. First, shallower waters tend to have higher temperatures, potentially allowing juveniles to achieve more rapid growth. Indeed, the preference of deep-sea larvae for shallower environments has been frequently reported (Yahagi et al., 2017; Gary et al., 2020). Second, intraspecific competition may drive smaller individuals away from deeper habitats. For example, in eastern Australian waters, multiple *Bathynomus* species coexist, with larger species occupying deeper habitats (Lowry and Dempsey, 2006). Similarly, west of Kyushu, limited food availability in the deep sea may induce competitive conspecific exclusion, contributing to the observed non-uniform size distribution.

In this study, sex identification and reproductive staging of *Bathynomus doederleini* were not conducted. Reproductive characteristics of this species have been studied in eastern Taiwan, where manca larvae (new recruits) were most abundant from March to September (Soong and Mok, 1991). Similarly, new recruitment of *Bathynomus giganteus* in the Gulf of Mexico has been reported primarily in winter and spring (Barradas-Ortiz et al., 2003). However, information about reproduction and brooding behavior of *B. doederleini* remains limited. This is likely due to the low probability of capturing brooding individuals using baited traps, as such individuals are not attracted to bait (Soong and Mok, 1991). In fact, at Japanese aquaria, brooding females kept in captivity took over a year from capture to hatching. This prolonged brooding period is exceptionally long among isopods. For example, the Antarctic isopod, *Ceratoserolis*, has a brooding period of approximately two years (Wägele, 1986), suggesting that *B. doederleini* may have similarly extended brooding durations as an adaptation to the cold deep-sea environment. Given the important ecological role of scavengers like *B. doederleini*, more comprehensive investigations are required to better understand its distribution and biology in the East China Sea. Future studies should incorporate both wide-area baited trap surveys and trawl sampling to capture individuals across a broader range of life stages.

## Acknowledgements

We are grateful to Tsunefumi Kobayashi, Toshiya Kawaguchi, Yurika Ono, Shoko Nishitsuji, and staff from the Fish and Ships Laboratory, Faculty of Fisheries, Nagasaki University, for supporting field studies. We also thank the crews of the training vessel Kakuyo-maru for providing the research cruise. Finally, we thank Steven D. Aird, Technical Editor (https://www.sda-technical-editor.org/), and the journal editors and anonymous reviewers for their valuable comments and suggestions that greatly improved the quality of the manuscript. This work was supported by JSPS KAKENHI (Grant Number: JP21K06337 to M.Y.), and the Sasakawa Scientific Research Grant from The Japan Science Society (Grant Number: 2024-4075 to S.A.).

## Author Contributions

YO Investigation, Writing – original draft, Visualization. ST Investigation, Writing – review & editing. KH Investigation, Writing – review & editing. TM Investigation, Writing – review & editing. SF Investigation, Writing – review & editing. KS Investigation, Writing – review & editing. MY Conceptualization, Methodology, Investigation, Writing – original draft, Supervision, Funding acquisition, Visualization.

## Supporting Information

Additional supporting information can be found online in the Supporting Information.

