## Supplementary Information for "Intraspecific differences in habitat depth in a deep-sea isopod, *Bathynomus doederleini* (Crustacea: Isopoda: Cirolanidae), off the west coast of Kyushu, Japan"

**Table S1** Summary of baited traps. The date, latitude, and longitude of each location, Depth (m), Water temperature (ºC), the number of traps that were undamaged at the time of retrieval, and trap time (hour), the time between trap set and retrieval.

| Voyage | Station | Date  (yy/m/d) | Latitude  (º-’, N) | Longitude  (º-’, E) | Depth (m) | Temperature  (ºC) | Rectangular trap | Cylindrical trap | Trap time  (h) |
| --- | --- | --- | --- | --- | --- | --- | --- | --- | --- |
| 1 | 5 | 2021/12/3 | 32-25.09 | 129-19.13 | 223 | 12.9 | 2 | 1 | 22.3 |
| 1 | 6 | 2021/12/3 | 32-14.74 | 129-19.20 | 278 | 11.8 | 2 | 1 | 20 |
| 1 | 7 | 2021/12/3 | 32-15.36 | 129-06.92 | 267 | 12.6 | 2 | 1 | 17.55 |
| 2 | 3 | 2022/5/13 | 32-28.77 | 129-07.54 | 253 | 12.2 | 2 | 0 | 24 |
| 2 | 4 | 2022/5/13 | 32-23.60 | 129-06.99 | 312 | 11 | 2 | 1 | 24 |
| 3 | 1 | 2022/10/19 | 32-40.26 | 129-06.75 | 151 | 16.7 | 2 | 1 | 18.78 |
| 3 | 2 | 2022/10/19 | 32-33.86 | 129-06.75 | 201 | 14.6 | 2 | 1 | 18.88 |
| 3 | 3 | 2022/10/19 | 32-28.86 | 129-06.65 | 249 | 12.6 | 2 | 1 | 19.08 |
| 3 | 4 | 2022/10/19 | 32-24.19 | 129-06.75 | 312 | 10.4 | 2 | 1 | 20.88 |
| 3 | 8 | 2022/10/21 | 32-19.15 | 129-03.92 | 337 | 9.9 | 2 | 1 | 23.08 |
| 3 | 9 | 2022/10/21 | 32-12.05 | 129-06.31 | 550 | 6.5 | 2 | 1 | 20.77 |
| 3 | 10 | 2022/10/21 | 32-08.28 | 129-06.15 | 591 | 6.4 | 2 | 1 | 19.71 |
| 4 | 4 | 2022/12/7 | 32-24.34 | 129-06.77 | 311 | 10.5 | 1 | 1 | 17.7 |
| 4 | 11 | 2022/12/8 | 31-55.88 | 129-21.88 | 739 | 5.8 | 2 | 1 | 20.58 |
| 4 | 12 | 2022/12/8 | 31-56.31 | 129-21.35 | 764 | 5.7 | 2 | 1 | 18.32 |
| 5 | 15 | 2024/4/26 | 32-07.24 | 129-16.48 | 542 | 7.2 | 2 | 1 | 18.35 |
| 5 | 14 | 2024/4/26 | 32-10.12 | 129-23.12 | 536 | 7.4 | 2 | 1 | 20.85 |
| 5 | 17 | 2024/4/26 | 32-13.12 | 129-30.00 | 448 | 7.8 | 1 | 1 | 16.67 |
| 6 | 17 | 2024/8/16 | 32-13.15 | 129-29.8 | 450 | 7.6 | 2 | 1 | 20.75 |
| 6 | 13 | 2024/8/16 | 32-06.00 | 129-40.8 | 652 | 7.6 | 2 | 1 | 17.65 |
| 6 | 16 | 2024/8/16 | 32-05.19 | 129-44.3 | 420 | 9.6 | 2 | 1 | 15.97 |
| 7 | 18 | 2024/10/17 | 32-04.77 | 129-27.4 | 644 | 5.8 | 2 | 1 | 19.62 |
| 7 | 19 | 2024/10/17 | 31-54.67 | 129-16.2 | 821 | 5.7 | 2 | 1 | 16.12 |

**Table S2** Complete list of captured organisms. Although not quantified, Amphipoda sp. and Ophiuroidea sp. were also collected at depths >500 m.

| Species ID | Order | Family | Species Name | Occurrence of Station | Numbers of individuals |
| --- | --- | --- | --- | --- | --- |
| *1* | Isopoda | [Cirolanidae](https://en.wikipedia.org/wiki/Cirolanidae) | *Bathynomus doederleini* | 2,3,4,5,6,7,8,9,10,11,12,13,14,15,16,17,18 | 1152 |
| *2* | Myxiniformes | Myxundae | *Myxine* sp. | 19 | 5 |
| *3* |  |  | *Eptatretus burger* | 1,4,15,17 | 45 |
| *4* |  |  | *Eptatretus okinoseanus* | 4 ,9,10,11,12,19 | 15 |
| *5* | Carcharhiniformes | Scyliohinidae | *Cephaloscyllium umbratile* | 3,5 | 3 |
| *6* |  |  | *Scyliorhinus torazame* | 3,5,6,7 | 17 |
| *7* | Dalatiidormes | Etmopteridae | *Etmopterus brachyurus* | 7 | 1 |
| *8* |  |  | *Etmopterus lucifer* | 18 | 1 |
| *9* | Anguulliformes | Synaphobranchidae | *Simenchelys parasitica* | 12,19 | 28 |
| *10* |  | Ophichthidae | *Ophichthus urolophus* | 2,3,4,5,6,7 | 61 |
| *11* | Gadiformes | Moridae | *Physiculus japonicus* | 4 | 1 |
| *12* | Ophidiiformes | Ophidiidae | *Neobythites sivicolus* | 12 | 1 |
| *13* | Scorpaeniformes | Sebastidae | *Helicolenus hilgendorfi* | 5,6,7,17 | 5 |
| *14* | Pleocymata | Scyliohinidae | *Ibacus ciliates* | 1 | 1 |
| *15* |  | Pandalidae | *Heterocarpus* sp. | 17 | 1 |
| *16* | Anomura | Digenidae | *Dardanus pedunculatus* | 1 | 1 |
| *17* |  | Galatheidae | sp. | 4 | 1 |
| *18* | Brachyura | Majidae | sp. | 4 | 1 |
| *19* |  | Oregoniidae | sp. | 3 | 1 |
| *20* |  | Goneplacidae | *Carcinoplax surgensis* | 2 | 14 |
| *21* |  | Epialtidae | *Chorilia japonica* | 13,18 | 3 |

**Table S3** Summary of baited traps with Species IDs.

| Voyage | Station | Rectangular trap of capture species ID | Cylindrical trap of capture species ID |
| --- | --- | --- | --- |
| 1 | 5 | 1,5,10,13 | 1 |
| 1 | 6 | 1,6,10,13 | 1 |
| 1 | 7 | 1,6,7,13 | 10 |
| 2 | 3 | 1,5,19 | - |
| 2 | 4 | 1,5,11,17,18 | 1 |
| 3 | 1 | 14,16 | 3 |
| 3 | 2 | 1,10, 20 | 1 |
| 3 | 3 | 6,10 | 1 |
| 3 | 4 | 1,3,10 | 1 |
| 3 | 8 | 1 | 1 |
| 3 | 9 | 1,4 | 1 |
| 3 | 10 | 1,4 | 1 |
| 4 | 4 | 1,3,5,10 | 1,3 |
| 4 | 11 | 1,4 | 1 |
| 4 | 12 | 1,4,9,12 | 9 |
| 5 | 15 | 1,3 | 1 |
| 5 | 14 | 1 | 1 |
| 5 | 17 | 1,3 | 15 |
| 6 | 17 | 1,13 | 1 |
| 6 | 13 | 1,21 | 1 |
| 6 | 16 | 1 | 1 |
| 7 | 18 | 1,8,21 | 1 |
| 7 | 19 | 2,4 | 2,9 |

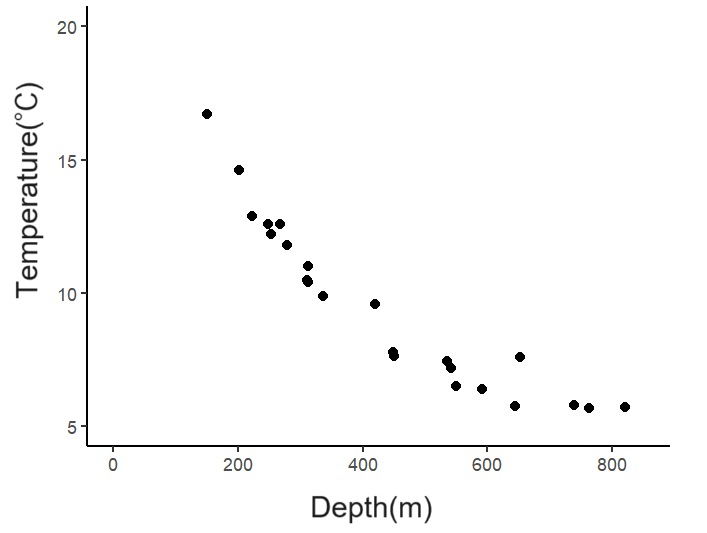

**Fig. S1** Relationship between water temperature and depth in the East China Sea (off the west coast of Kyushu, Japan).
